# *Streptococcus pneumoniae* modulates reassortment of Influenza A Virus in a Pneumolysin dependent manner

**DOI:** 10.64898/2026.02.02.703233

**Authors:** Hannah M. Rowe

**Affiliations:** Department of Microbiology, Oregon State University, Corvallis OR USA

## Abstract

Influenza A viruses (IAV) can undergo rapid evolution by acquisition of new genes through reassortment between IAV strains leading to immune evasion, antiviral resistance, and change in host range. Understanding of the factors that can enhance or reduce reassortment frequency therefore has implications for protection of individual and public health. Reassortment requires co-infection by two or more viral particles to the same host cell. Prior studies had identified the potential for IAV particles to aggregate on the bacterial surface of *Streptococcus pneumoniae* and other respiratory pathobionts, suggesting that bacterial cells could serve as a method to aggregate IAV particles and potentially facilitate reassortment. To test this hypothesis, a temperature sensitive, oseltamivir resistant viral strain was generated and used in *in vivo* and *in vitro* co-infection experiments with wild type virus in the presence of live and killed bacterial cells. While killed pneumococcal cells can enhance IAV reassortment frequency in a density-dependent manner, live pneumococci cannot, suggesting a product produced by viable pneumococci inhibits viral reassortment. Genetic deletion of the pneumococcal cholesterol-dependent cytolysin, Pneumolysin (Ply), in combination with inhibition of inflammatory signaling induced by Ply, restored the enhancement of reassortment to the levels seen with killed pneumococci. The results of this work suggest that the bacterial cells that colonize the human upper respiratory tract can have a complex role in modulating IAV reassortment frequency. Bacterial cells are capable of facilitating enhancement of viral reassortment, however, bacterial products can negate these effects. This work demonstrates the role of one such interaction with Ply inhibiting the enhanced reassortment otherwise conferred by *S. pneumoniae* cells. Overall, this suggests a new model whereby the human nasopharyngeal microbial community which differs from individual to individual and across the lifespan can impact IAV evolutionary dynamics, with some bacterial cells and metabolites increasing and others decreasing IAV reassortment frequency.

**Summary:** Influenza A virus can evolve rapidly by exchanging genes between viral strains, leading to vaccine failure, spread of antiviral resistance, and potential pandemic emergence. Bacterial cells may be able to increase the frequency of viral genetic exchange through concentration of viral particles on the bacterial cell surface. Prior findings had shown that *Streptococcus pneumoniae,* a frequent colonizer of the human upper respiratory tract, can directly bind to IAV particles. This work showed in cell culture and animal models that killed *S. pneumoniae* cells can increase viral genetic exchange. However, live bacteria capable of producing the bacterial toxin Pneumolysin are unable to promote viral genetic exchange. This suggests that bacterial modulation of viral evolutionary dynamics is a complex interaction whereby bacterial cells and products can both enhance and reduce viral genetic exchange, and the human nasopharyngeal microbial community which differs from person to person and on human age could have a role in Influenza A virus evolutionary dynamics.

## Introduction

Influenza A virus (IAV) possesses a segmented genome, which facilitates rapid genomic change through reassortment of genome segments during co-infection of a single host with two or more IAV strains. Each major human pandemic of the 20^th^ and 21^st^ century has resulted from a reassortment event(1–3). Novel avian IAV strains are also generated through reassortment events leading to panzootics, including the ongoing H5 outbreak(4–6), and the recent host range expansion to cattle(7). While these consequences are the most dramatic, reassortment can also lead to spread of antiviral resistance genes(8, 9) with resultant impacts on infection prevention and treatment options.

The frequency of reassortment has been historically underappreciated but does occur at high levels during natural infections and in cell culture and animal models(10–15). Most reassortant strains though have reduced infection or transmission fitness(10). However, the rare increased fitness resulting from a reassortment event has high consequences for individual and public health, as well as economic productivity and food security. Therefore it is critical to understand the mechanisms responsible for increased reassortment frequency.

One recently described mechanism for enhancement of viral reassortment is aggregation of viral particles, as seen in Reoviruses(16). Direct interactions between IAV particles and bacterial cells have recently been described(17–19), demonstrating that multiple viral particles on the surface of a single bacterial cell. This suggests a model whereby a bacterial cell is able to aggregate viral particles. If those viral particles have different genetic composition, this could allow enhanced reassortment frequency through enhanced frequency of co-infection. Additionally, bacterial components or bacterial cells could alter host cell intracellular trafficking and therefore reassortment through altered trafficking of viral genome segments. Bacterially-enhanced viral horizontal gene transfer has been previously demonstrated with Poliovirus(20) and Adenovirus(21). In the case of Poliovirus, the identified mechanism was increased co-infection of host cells, with bacterial species that bind multiple viral particles better able to support recombination than bacterial species that bind single or no viral particles(20). In the case of Adenovirus the identified mechanism was the bacterial RecA enzyme, recognizing Chi sites within the viral genome, and directly promoting recombination(21).

Utilizing an oseltamivir resistant, temperature sensitive IAV strain in combination with a wild type (oseltamivir sensitive, temperature resistant) strain and measuring the output of temperature resistant, oseltamivir resistant strains following infection of *in vitro* and in murine models it appears that *Streptococcus pneumoniae* and other upper respiratory pathobionts are able to support enhanced reassortment of IAV. These data support the model of enhanced co-infection, as there is dose dependent response, with higher numbers of bacterial cells leading to higher output titer of oseltamivir resistant, temperature resistant virus. However, viability and subsequent pneumococcal toxin production reduce reassortment frequency, suggesting an additional role for bacterial metabolites to alter host cell responses and intracellular trafficking.

## Results

Prior work had identified the capacity for bacterial upper respiratory pathobionts to directly interact with viral particles, with multiple viral particles visualized on a single, or aggregate, of bacterial cells(17, 18). This suggested a potential mechanism for upper respiratory pathobionts to promote IAV reassortment. Using inverse PCR and mutagenic primers, temperature sensitive(22) PA and PB1 genes were generated, as was an oseltamivir resistant NA gene(23). HEK-293T cells were transfected with the three mutant plasmids and the remining five wild type plasmids to generate a temperature sensitive, oseltamivir resistant strain of A/Puerto Rico/08/1934 (H1N1) (PR8) to be utilized in this study.

To test the ability of *S. pneumoniae* cells to promote viral reassortment *in vivo,* mice were treated with oseltamivir and infected with combinations of wild type and temperature sensitive drug resistant virus and *S. pneumoniae*. The mouse core body temperature will only support replication of wild type virus, or of reassortant virus that has gained the wild type polymerase, while the oseltamivir treatment will select for virus that has the resistant neuraminidase. Equal quantities of wild type and temperature sensitive/oseltamivir resistant virus, as determined by 50% tissue culture infectious dose (TCID_50_), were mixed with live or ethanol-fixed *S. pneumoniae*, as previous studies(17) showed that both viable and ethanol killed pneumococci are equally capable of binding IAV particles. Bacterial cells and associated viral particles were pelleted by low speed centrifugation. Assuming a 1% input virus bound, as previously demonstrated(17), control inocula were prepared with one hundred fold less virus for each virus alone and a for a mix of both viruses. To select for reassortant viruses with the oseltamivir resistant neuraminidase, mice were treated with 25mg/kg oseltamivir phosphate by oral gavage two hours prior to viral infection and every twelve hours post viral challenge. Live *S. pneumoniae* would be capable of colonizing and supporting multiple potential rounds of reassortment in the respiratory tract as progeny viruses associated with colonizing bacteria. Ethanol killed *S. pneumoniae* would be cleared from the respiratory tract but would support delivery of multiple viral particles to the same host cell in the first round of replication. Mice were weighed daily for four days to determine signs of severe viral illness. Weight loss was only observed in mice infected with co-sedimented bacterial-viral complexes (Figure 1A), suggesting that bacterial co-administration was promoting viral reassortment to allow viral replication in the presence of drug pressures which would not allow replication of wild type virus. To verify the presence of reassortant virus, at 4 days post infection lungs were harvested and both total viral load and oseltamivir resistant temperature resistant viral load was determined by TCID_50_ in the presence and absence of oseltamivir. Temperature resistant, oseltamivir resistant virus was only recovered from mice infected with co-sedimented bacterial-viral complexes (Figure 1B) supporting bacterially mediated reassortment.

**Figure 1:**
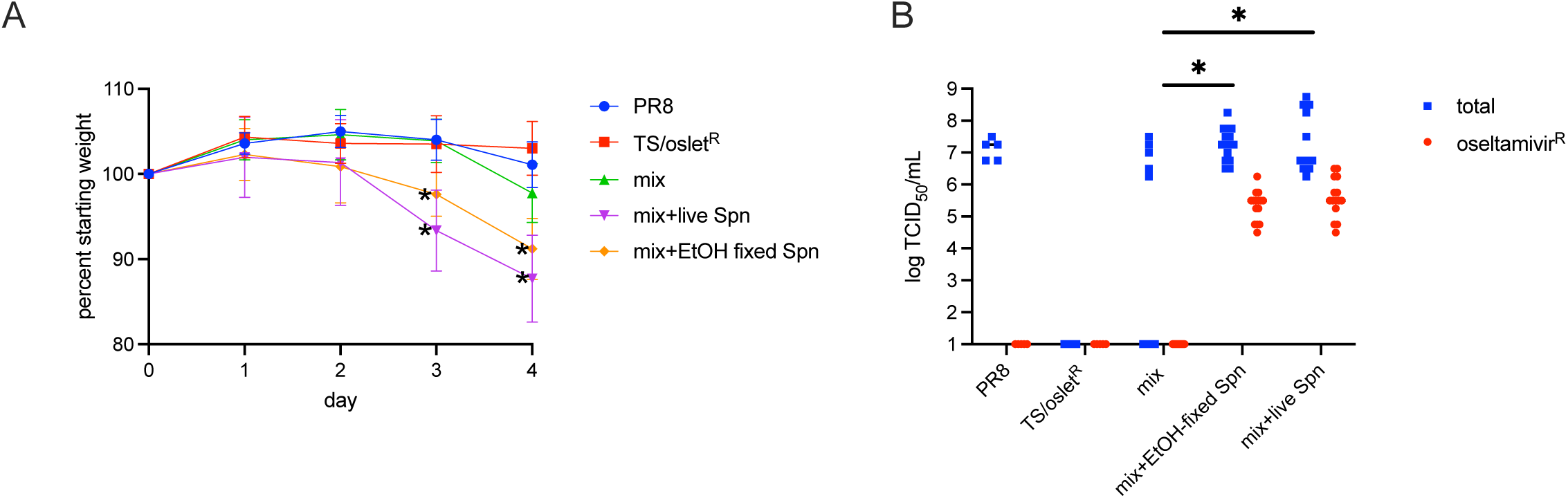
Co-sedimentation with *S. pneumoniae* increases viral reassortment *in vivo.* Mice were infected with wild type A/Puerto Rico/08/1934 (H1N1) (PR8), a temperature sensitive and oseltamivir resistant derivative of PR8 (TS/oslet^R^), a mix of both viruses (mix), a mix of both viruses co-sedimented with live *S. pneumoniae* (mix+live Spn), or a mix of both viruses co-sedimented with ethanol-fixed *S. pneumoniae* (mix+EtOH fixed Spn), and were treated with 25mg/kg oseltamivir phosphate at 2 hours prior to infection and every 12 hours post infection for 4 days. **A)** Each mouse was weighed daily post infection and weight was compared to pre-infection weight. Data are presented as mean with error bars representing standard deviation. *= p<0.01 by unpaired t-test compared to mice infected with mix of viruses on that date post infection. **B)** Viral titers in lungs of mice 4 days post infection. Total titer (blue squares) and oseltamivir resistant titer (red circles) was determined by TCID_50_ in the absence and presence of 23μM oseltamivir carboxylate. Data are presented as titer from each animal is a single data point with the bar representing the median. * = p<0.01 by Mann Whitney. 5-15 mice per infection condition were used in 2 independent infection replicates.

### Impact of pneumococcal density and viability on viral reassortment

To determine the roles of both the concentration and the viability of bacterial cells on viral reassortment frequency, equal quantities (4 log TCID_50_/mL) of wild type and temperature sensitive, oseltamivir resistant virus were pre-mixed with 10^5^, 10^6^, or 10^7^ CFU/mL of live or ethanol killed *S. pneumoniae* in infection media for 30 minutes prior to incubation. Cells were infected for one hour, followed by removal of inoculum and replacement with infection media supplemented with antibiotics for a one-step viral growth curve to measure the burst size of temperature resistant, oseltamivir resistant virus progeny. Therefore, live bacteria were only present during the formation of the bacterial-viral complex, and during initial viral attachment and entry, not for the entire viral lifecycle. To determine if upper and lower respiratory cells behave similarly for supporting reassortment, both A549 lung adenocarcinoma, and Detroit 562 nasopharyngeal carcinoma cell lines were used. For both A549 (Figure 2A) and Detroit 562 (Figure 2B) increasing the amount of ethanol killed bacteria generally increased the recovery of temperature resistant, oseltamivir resistant (probable reassortant) viral progeny. This supports the hypothesis that increased bacterial count allows increased delivery of multiple viruses to the same cell. However, when using live *S. pneumoniae* the same trend was not observed. The presence of live bacteria reduced the recovery of temperature resistant, oseltamivir resistant virus, with the lowest concentration of bacteria having the least impact. This supports a second hypothesis that there is something produced by live *S. pneumoniae* that inhibits reassortment.

**Figure 2:**
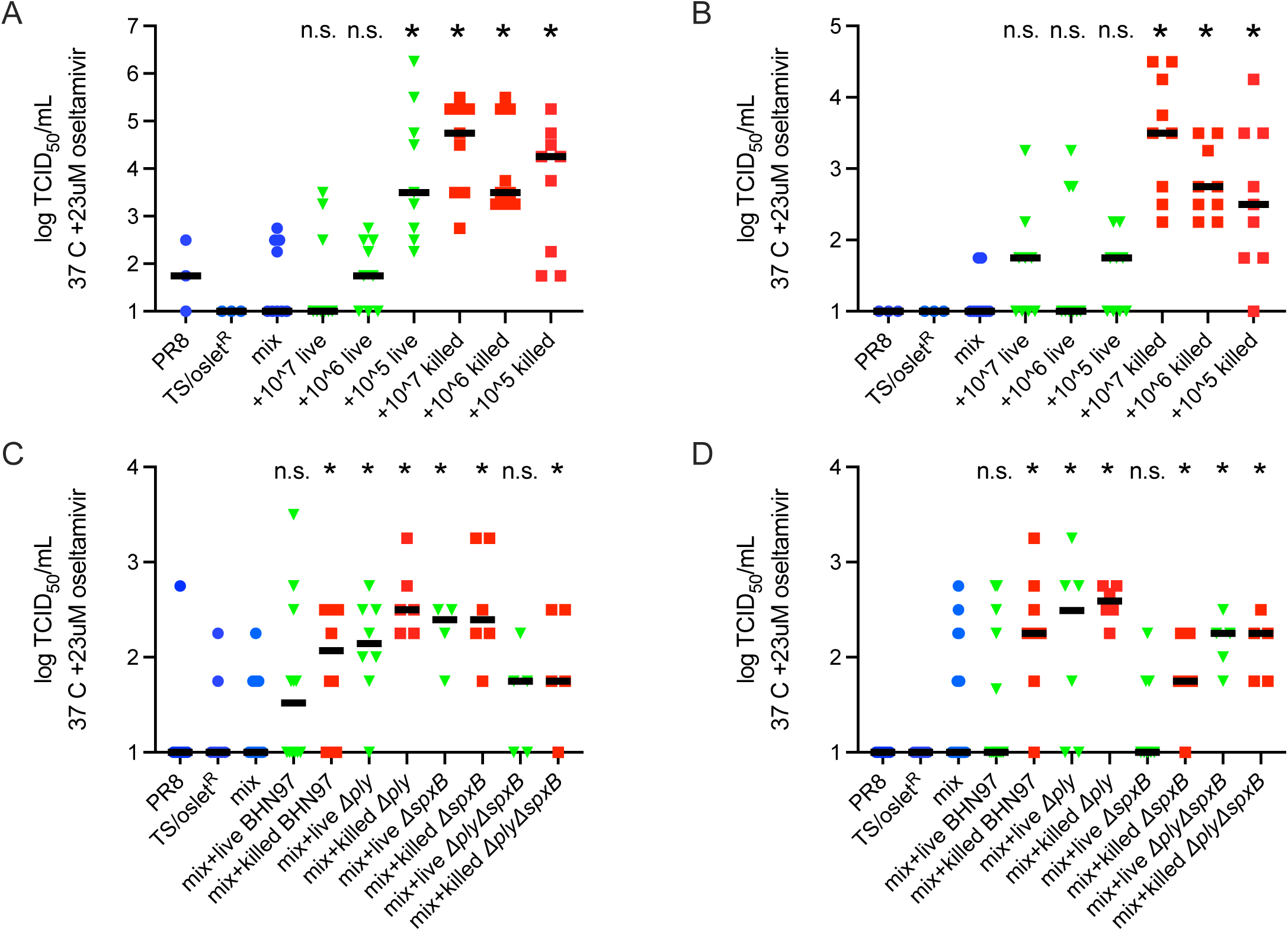
Viral reassortment is modulated by concentration of pneumococcal cells and by toxic products produced by viable *S. pneumoniae*. A549 (**A** and **C**) or Detroit 562 cells (**B** and **D**) were infected with wild type A/Puerto Rico/08/1934 (H1N1) (PR8), a temperature sensitive and oseltamivir resistant derivative of PR8 (TS/oslet^R^), a mix of both viruses (mix), a mix of both viruses co-sedimented with live or ethanol-fixed *S. pneumoniae* at indicated concentrations (**A** and **B**), or 10^6^ CFU/mL wild type strain BHN97 or isogenic mutants (**C** and **D**). Following one hour of incubation, inocula were removed and replaced with viral infection media containing 10X working concentration of penicillin/streptomycin, cells were incubated for 24 hours at permissive conditions (33 °C and without oseltamivir). Reassortant progeny viruses were determined by titration on MDCK cells at non-permissive conditions (37 °C with 23μM oseltamivir carboxylate). Data are presented as titer from each cell culture well as a single data point with the bar representing the median. n.s.= not significant, * = p<0.01 by Mann Whitney compared to mix of viruses without co-sedimented bacteria. 3-9 wells per infection condition from 3 independent infection replicates.

### Impact of pneumococcal toxin and peroxide production on viral reassortment

Two potential candidate bacterial metabolites were examined due to their ability to act as damaging agents to mammalian cells. *S. pneumoniae* produces a cholesterol-dependent cytolysin, pneumolysin (Ply), and millimolar(24, 25) quantities of hydrogen peroxide during central metabolism, mediated by pyruvate oxidase, SpxB(24, 26). To determine the roles of these bacterial products, mutants in *spxB*, *ply,* or double mutants were used in the assays described above with a single bacterial concentration (10^6^ CFU/mL). As shown above, infection of A549 (Figure 2C) or Detroit 562 (Figure 2D) cells with live wild type *S. pneumoniae* did not promote reassortment significantly above co-infection with a mix of viruses alone, while infection with ethanol-killed wild type bacteria did. Both live and ethanol-killed *Δply* bacteria were able to promote reassortment in both A549 (Figure 2C) and Detroit 562 (Figure 2D) cells, suggesting that Ply could be a factor released by viable pneumococci responsible for inhibition of reassortment. With regards to hydrogen peroxide production, ethanol-killed *ΔspxB* bacteria were able to promote reassortment in both A549 and Detroit 562 cells, while live *ΔspxB* cells were able to promote reassortment in A549 cells (Figure 2C), but not Detroit 562 cells (Figure 2D) suggesting a difference between upper and lower respiratory cells. While reducing bacterial H_2_O_2_ production was sufficient to allow reassortment in the presence of viable bacteria in A549 cells, it was not sufficient in Detroit 562 cells, suggesting an additional factor produced by viable pneumococci is still inhibiting viral reassortment in upper-respiratory like cells. When the viral strains were mixed with a bacterial strain with both *ply* and *spxB* deleted, ethanol-killed *ΔplyΔspxB* bacteria were able to promote reassortment in both A549 and Detroit 562 cells, whereas live *ΔplyΔspxB* were only able to promote reassortment in Detroit 562 cells (Figure 2 C and D).

Notably, neither the presence of viable or killed *S. pneumoniae,* nor the expression of *spxB* nor *ply*, impacts replication of wild type IAV strain PR8 in A549 or Detroit 562 cells (supplemental figure 1A and 1B). This suggests that ethanol-killed bacteria do not enhance reassortment through merely increasing viral replication, rather that the co-infection in presence of killed bacterial cells has an impact on reassortment specifically. Additionally this supports a hypothesis that *ply* and *spxB* expression by *S.pneumoniae* does not inhibit viral replication overall, rather acts specifically on viral reassortment.

### TLR4 antagonism, but not peroxide neutralization by catalase reverses pneumococcal mediated reduction of reassortment

As SpxB activity can also be important for release of Ply(27), to isolate the role of hydrogen peroxide generation, co-infections were performed similar to as above, with catalase to neutralize bacterially-produced H_2_O_2_. Catalase was present during the pre-incubation of bacterial cells with viral particles, during the initial attachment of viral particles, and during the one-step viral growth curve. Incubation with catalase did not increase viral reassortment frequency with or without bacterial co-infection in either A459 (Figure 3A) or Detroit 562 (Figure 3B) cells. This suggests that it is more likely that Ply is the factor produced by viable pneumococci that reduces viral reassortment. However, it is possible that other structural and metabolic differences(28–30) present in *spxB* deletion mutant bacteria are also responsible.

**Figure 3:**
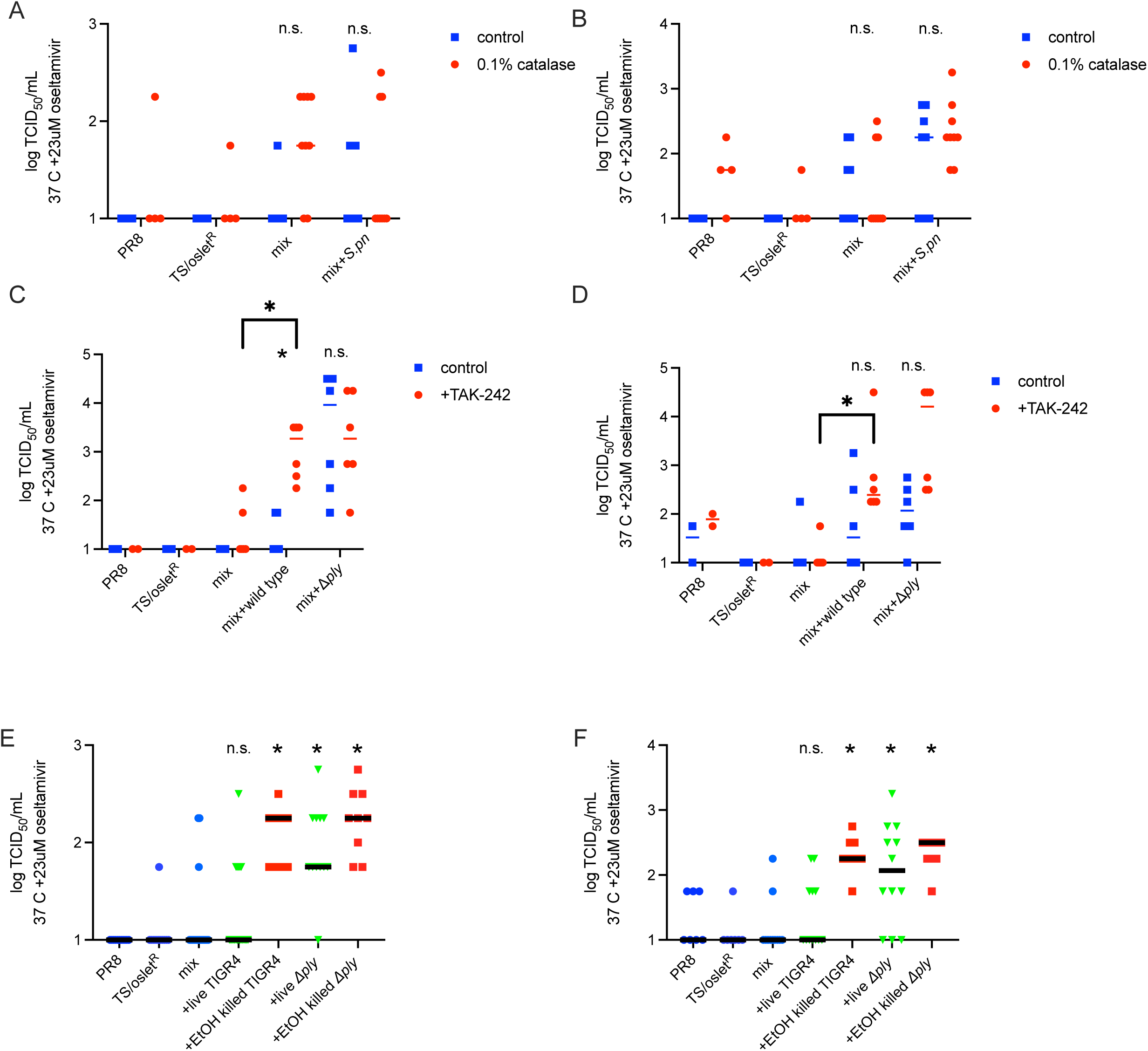
Modulation of viral reassortment by *S. pneumoniae* is dependent on pneumolysin production by viable pneumococci. A549 (**A**, **C** and **E**) or Detroit 562 cells (**B**, **D** and **F**) were infected with wild type A/Puerto Rico/08/1934 (H1N1) (PR8), a temperature sensitive and oseltamivir resistant derivative of PR8 (TS/oslet^R^), a mix of both viruses (mix), a mix of both viruses co-sedimented with live or ethanol-fixed *S. pneumoniae*: (**A** and **B**) inocula were prepared and cells were infected in the presence or absence of 0.1% catalase to determine the role of *spxB* mediated H_2_O_2_ production on viral reassortment. (**C** and **D**) cells were pre-treated with, inocula were prepared in, and cells were infected in the presence of 1uM TAK-242, or 0.004% DMSO vehicle control. (**E** and **F**) *S. pneumoniae* strain TIGR4 and isogenic *ply* deletion mutant were used to determine bacterial strain dependence of *ply* interference in viral reassortment. Following one hour of incubation, inocula were removed and replaced with viral infection media containing 10X working concentration of penicillin/streptomycin, cells were incubated for 24 hours at permissive conditions (33 °C and without oseltamivir). Reassortant progeny viruses were determined by titration on MDCK cells at non-permissive conditions (37 °C with 23μM oseltamivir carboxylate). Data are presented as titer from each cell culture well as a single data point with the bar representing the median. n.s.= not significant, * = p<0.01 by Mann Whitney compared to mix of viruses without co-sedimented bacteria. 3-21 wells per infection condition from at least 3 independent infection replicates.

In addition to its role as a cholesterol dependent cytolysin causing direct lysis of mammalian cells, Ply is also immunostimulatory through its role as a Toll-like receptor 4 (TLR4) agonist(31). To investigate the role of TLR4 signaling on bacterial mediated viral reassortment, the TLR4 inhibitor TAK242(32) was utilized during co-infection experiments. Cells were pre-treated for 24 hours prior to infection, and TAK242 or vehicle control was present during pre-incubation of bacterial cells with viral particles, during the initial attachment of viral particles, and during the one-step viral growth curve. For both A549 (Figure 3C) and Detroit 562 (Figure 3D), if TLR4 was inhibited with TAK242, live wild type bacteria were now able to increase the reassortment frequency when compared to a mix of the two virus strains without bacterial co-incubation. In A549 cells (Figure 3C) TAK242 treatment significantly improved reassortant recovery when viruses were co-infected with live wild type bacteria compared to viruses co-infected with live wild type bacteria treated with vehicle control, while for Detroit 562 cells (Figure 3D) there is no significant difference between the TAK242 and vehicle control groups when co-infected with live wild type bacteria, suggesting a potential cell type dependence difference in Ply signaling or TLR4 response. The difference in reassortant output was the same between TAK242 and vehicle treatment for when incubated with *Δply* bacteria for both A549 (Figure 3C) and Detroit 562 (Figure 3D) cells. This suggests that Ply action through TLR4 is altering reassortment. To further support the role of *ply* in reducing viral reassortment output, a *Δply* mutant generated in a different strain of *S. pneumoniae*, TIGR4, was utilized in co-infection and reassortment experiments. As in strain BHN97, ethanol-killed TIGR4 enhances viral reassortment, but live TIGR4 reduces reassortment in a *ply* dependent manner in both A549 and Detroit 562 cells (Figure 3E and 3F).

Notably, this effect is specific to reassortment, and not to general viral replication alteration in the presence of TLR4 inhibition. For neither A549 (Supplemental Figure 2A) nor Detroit 562 (Supplemental Figure 2B) co-infection of bacteria with wild type IAV strain PR8 does treatment with TAK242 alter viral replication, in the presence of live wild type or *Δply* bacteria. This suggests that the TLR4 signaling mediated by Ply is specifically reducing reassortment, not viral replication overall.

### Other respiratory pathobionts

To determine if the alteration in viral reassortment is specific to *S. pneumoniae* or common to other respiratory pathobionts, the co-infection experiments were done with three other upper-respiratory pathobionts: non-typable *Hemophilus influenzae* (Figure 4A)*, Moraxella catarrhalis* (Figure 4B) and *Staphylococcus aureus* (Figure 4C). To determine the roles of both the concentration and the viability of bacterial cells on viral reassortment frequency, equal quantities (4 log TCID_50_/mL) of wild type and temperature sensitive oseltamivir resistant virus were pre-mixed with 10^5^, 10^6^, or 10^7^ CFU/mL of live or ethanol killed bacteria followed by infection of A549 cells and assessment of reassortant virus progeny after a one step viral growth curve. For *H. influenzae*, ethanol-killed bacteria were able to promote reassortment, while live bacteria could not (Figure 4A), and for *M. catarrhalis,* only the highest concentration of killed bacteria and no concentration of live bacteria could promote reassortment (Figure 4B), but for *S. aureus* the opposite trend was seen where live bacteria supported reassortment, while ethanol-killed bacteria could not (Figure 4C). This suggests that not all ethanol-fixed bacterial cells can serve to facilitate viral co-infection and therefore reassortment. It also suggests that a factor produced by viable *H. influenzae* inhibits reassortment, while a factor produced by viable *S. aureus* enhances viral reassortment.

**Figure 4:**
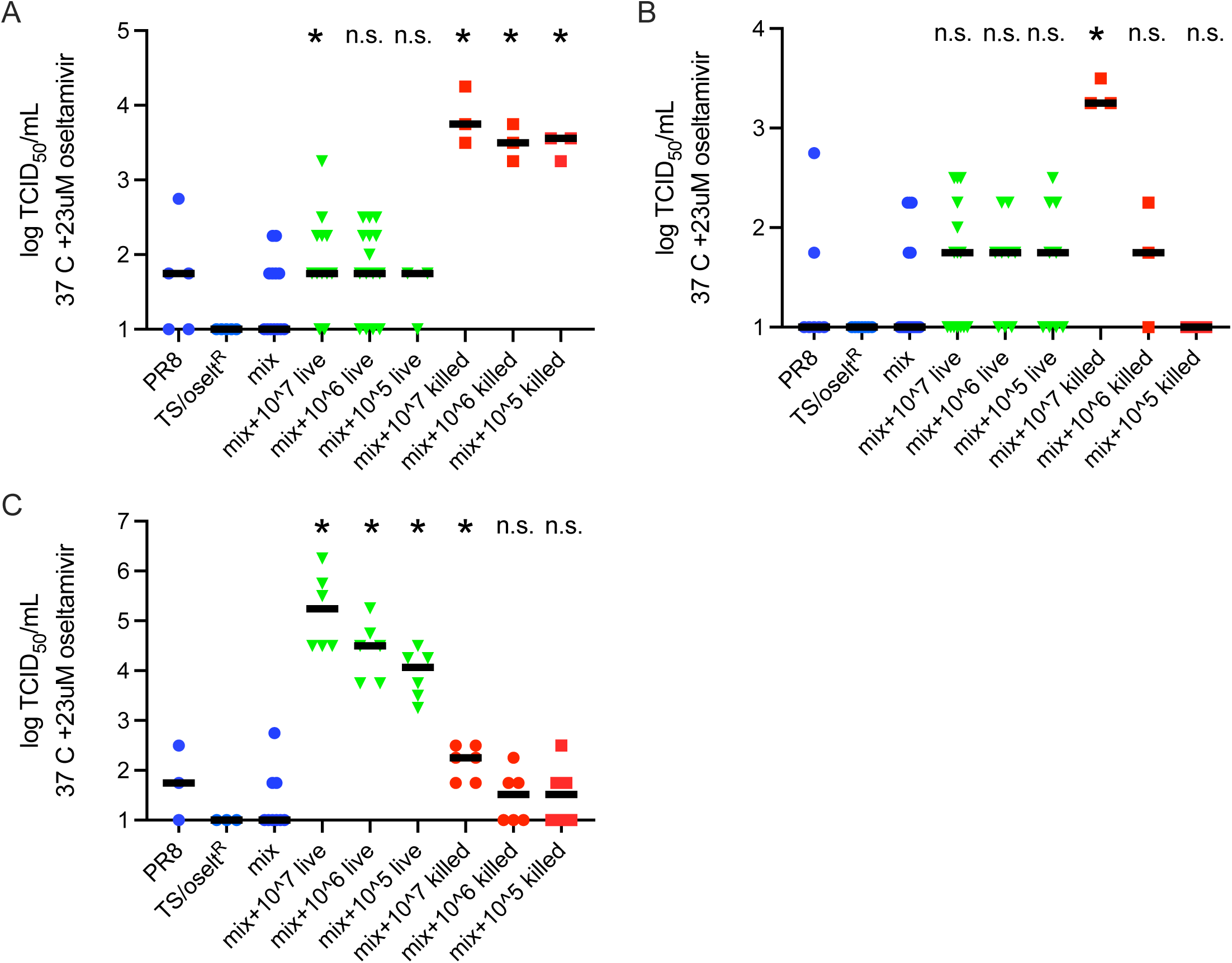
Other respiratory pathobionts also modulate viral reassortment in a density or viability dependent manner. A549 cells were infected with wild type A/Puerto Rico/08/1934 (H1N1) (PR8), a temperature sensitive and oseltamivir resistant derivative of PR8 (TS/oslet^R^), a mix of both viruses (mix), a mix of both viruses co-sedimented with indicated concentration of live or ethanol-fixed (**A**) Non-typable *H. influenzae*, (**B**) *M. catarrhalis*, (**C**) *S. aureus*. Following one hour of incubation, inocula were removed and replaced with viral infection media containing 10X working concentration of penicillin/streptomycin, cells were incubated for 24 hours at permissive conditions (33 °C and without oseltamivir). Reassortant progeny viruses were determined by titration on MDCK cells at non-permissive conditions (37 °C with 23μM oseltamivir carboxylate). Data are presented as titer from each cell culture well as a single data point with the bar representing the median. n.s.= not significant, * = p<0.01 by Mann Whitney compared to mix of viruses without co-sedimented bacteria. 3-16 wells per infection condition from at least 3 independent infection replicates.

## Discussion

Overall, this study suggests a model whereby bacterial cells and their metabolites can act directly on, or indirectly through immunomodulation of, host cells to alter IAV reassortment. This suggests that the microbial community that differs between individuals(33, 34), upon acute respiratory infection(35–38) and across the lifespan(39–41) may alter IAV evolutionary dynamics, in addition to the impacts previously demonstrated on transmission dynamics(42, 43).

This report expands our understanding of the well-characterized model of synergy between IAV and *S. pneumoniae* suggesting a new role of bacterial cells during viral infection and evolution, in addition to what has been previously demonstrated regarding the synergy in pathogenesis(44, 45). Utilizing an *in vivo* infection model, with an intact normal nasopharyngeal bacterial community, enhanced viral reassortment was seen when mice were infected with viral inocula pre-incubated with wild type or ethanol-fixed *S. pneumoniae*. Utilizing *in vitro* models of upper and lower respiratory cell lines, a density dependent increase in viral reassortment was seen with increasing concentrations of ethanol-fixed pneumococci. However, viable pneumococci were shown to not enhance viral reassortment. The results presented here show divergent results between the *in vivo* and *in vitro* ability of live *S. pneumoniae* able to support enhanced reassortment. These results could be explained by the theory that during *in vivo* infection, immune mediated and interbacterial competition mediated killing of pneumococcal cells, and therefore increased presence of dead pneumococcal cells able to act as sites to concentrate virions and enhance viral co-infection and therefore reassortment, or that the presence of pneumococci altered the natural murine nasopharyngeal community or the immune milieu to enhance reassortment. In *in vitro* models, genetic knockout of production of *spxB*, a major generator of hydrogen peroxide(24–26), or of ply, responsible for production of the cholesterol dependent cytolysin: Pneumolysin, restoration of enhanced reassortment was demonstrated, suggesting that the toxic or immunomodulatory effects of these pneumococcal metabolites was interfering with IAV reassortment. Notably, this impact on reassortment is independent of the impact on viral replication. The failure of neutralization of hydrogen peroxide to restore enhancement of reassortment in the presence of viable bacteria, in combination with the blockade of TLR4 signaling enhancing viral reassortment in the presence of viable bacteria suggests that pneumolysin, not hydrogen peroxide, is responsible for modulation of IAV reassortment. Finally, the impact of other common respiratory pathobionts was tested to determine if the modulation of IAV reassortment was specific to *S. pneumoniae*. *H. influenzae, M. catarrhalis* and *S. aureus* can enhance IAV reassortment to varying extents, and with differing dependence on bacterial viability.

Together these data suggest a complex role played by the nasopharyngeal microbial community and co-infecting bacterial pathogens during IAV infection, where viral particles can be concentrated on the surface of bacterial cells promoting co-infection and reassortment, as has been previously suggested and demonstrated for other viruses(20, 21). However, it also demonstrated that products produced by viable bacterial cells can negate this enhanced reassortment. This report suggests this can be done by the pneumococcal cholesterol dependent cytolysin via indirect action through signaling mediated by TLR4, though the impact of direct action cannot be excluded at this time. Other bacterial products could likewise enhance or reduce viral reassortment through direct and indirect activity, and the balance of reassortment-promoting and reassortment-reducing bacterial actions in an individual or population will depend on the constituents of the nasopharyngeal microbial community. This highlights the importance of characterizing the constituents and the functions provided by the nasopharyngeal microbiome and co-infecting bacterial pathogens to better understand and find ways to control IAV infection, transmission, and evolution.

## Materials and Methods

### Cell Culture

A549 (ATCC CCL-185) was cultured in F12-K media (Gibco) supplemented with 10% fetal bovine serum (Hyclone). Detroit 562 (ATCC CCL-138) was cultured in Minimal Essential Media (Gibco) supplemented with 10% fetal bovine serum (Hyclone). Madin-Darby Canine Kidney (MDCK) cells (ATCC CCL-34) in Minimal Essential Media (Gibco) supplemented with: GlutaMax (Gibco), Sodium Pyruvate (Gibco) and 10% fetal bovine serum (Hyclone). All cells were grown in a humidified incubator at 37 C with 5% CO_2_ atmosphere.

### Bacterial culture

*S. pneumoniae* strain BHN97 (Serotype 19F) or TIGR4 (Serotype 4) was grown on Tryptic Soy Agar (BD) supplemented with 3% sheep red blood cells (Lampire) in a humidified incubator at 37 C with 5% CO_2_ atmosphere. Solid media was supplemented with 1ug/mL erythromycin (Fisher) or 150 ug/mL spectinomycin (Sigma Aldrich) when selection of antibiotic resistant allelic replacement was required. Overnight growth was transferred to Todd Hewitt media (BD) supplemented with 0.2% yeast extract (BD) (THY media) in a humidified incubator at 37 C with 5% CO_2_ atmosphere. BHN97 *ΔspxB* deletion strain(46) and BHN97 *Δply* deletion strain(47) were generated in previous studies and TIGR4 *Δply* deletion strain was a gift from Elaine Tuomanen. The double *ΔspxB Δply* double deletion was generated by allelic exchange with an spectinomycin resistance cassette to delete *spxB* in a *ply* deletion background. Approximately 1 kb upstream and downstream of *spxB* was amplified with primers (see table 1) incorporating an overlap of the spectinomycin resistance cassette using PrimeStar High Fidelity Polymerase (Takara). PCR products were gel extracted (Qiagen MinElute) and along with the spectinomycin resistance cassette were combined using splicing by overlap extension PCR using PrimeStar High Fidelity Polymerase (Takara). Purified PCR product was transformed into BHN97*Δply* in THY media supplemented with 0.002% BSA, 0.2% glucose and 0.0002% CaCl_2_(48) grown to early log (OD_620_ approximately 0.2), diluted 1:100 in supplemented THY, incubated 14 minutes with 2μg/mL CSP-1 and CSP-2(49). Gene deletion was confirmed by absence of amplification with internal primers.

**Table 1:**
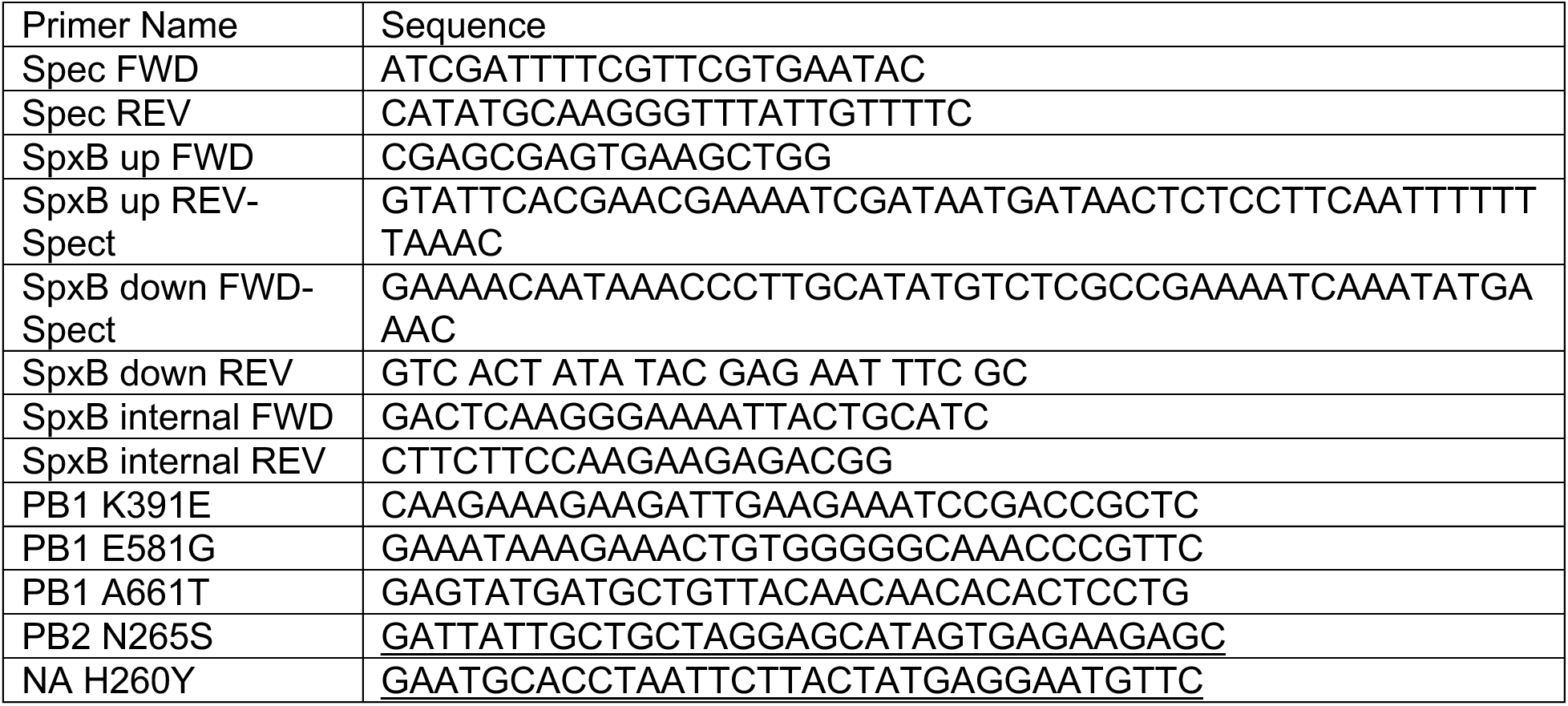
Primers used in this study.

The nontypeable *H. influenzae* strain 86-028NP was grown on chocolate agar supplemented with 11,000 units/L bacitracin and then directly inoculated into Brain Heart Infusion (BD) broth supplemented with 0.2% yeast extract, 10 μg/mL hemin, and 10 μg/mL NAD and grown with aeration to mid-log phase (50). *Staphylococcus aureus* strain MW2 and *Moraxella catarrhalis* (51) were grown on unsupplemented TSA plates, directly inoculated in Brain Heart Infusion broth supplemented with 0.2% yeast extract, and grown with aeration to mid-log for use in experiments.

### Ethanol killed bacteria

10^8^ CFU bacteria were resuspended in ice cold 70% ethanol and incubated on ice for 5 minutes. Cells were centrifuged at 10,000 xg and resuspended in 1mL phosphate buffered saline (PBS) (Gibco) or viral infection media depending on downward experiment. Loss of viability was determined by plating 100uL killed bacteria on a TSA/sheep blood plate and absence of growth verified.

### Viral culture

Influenza virus strain A/Puerto Rico/08/1934 (H1N1) (PR8) was grown in MDCK cells. Prior to infection cells were washed twice with PBS. Cells were infected at an MOI∼0.01 in Minimal Essential Media (Gibco) supplemented with: BSA fraction V (Gibco), GlutaMax (Gibco), and 1ug/mL TPCK-Trypsin (ThermoScientific). After 96 hours of infection and complete destruction of the monolayer, supernatant was harvested, centrifuged at 5000 xg to pellet debris and was aliquoted and stored at −70C. Titer was determined by 50% tissue culture infectious dose (TCID_50_) using the method of Reed and Munch(52).

#### Generation of Temperature Sensitive Oseltamivir Resistant Virus

The temperature sensitive oseltamivir resistant virus was generated using reverse genetics as described previously(53). For temperature sensitivity: K391E, E581G, and A661T substitutions were introduced into PB1 and N265S substitution was introduced into PB2(22), for oseltamivir resistance an H260Y (equivalent to H274Y(54)) substitution was introduced into NA(23). Mutations were inserted into the plasmids using the QuikChange Multi Site-Directed Mutagenesis Kit (Agilent) following manufacturer directions and primers in table 1. Successful mutation was confirmed with Sanger sequencing.

HEK-293T cells (ATCC CRL-11268) were plated in 6 well cell culture dishes (Costar) at 2×10^5^ cells per mL (∼80% confluency). Immediately prior to transfection, cells were washed and media replaced with OptiMEM (Gibco). HEK-293T cells were transfected with 1ug each of the 8 plasmids containing the influenza genome. Plasmids were diluted with OptiMEM and mixed with TransIT-LT1 Reagent (Mirus) according to manufacturer directions and incubated for 15 minutes at room temperature. TransIT-LT1:DNA complexes were added to cells and incubated for 24 hours at 33 degrees in a humidified 5% CO_2_ atmosphere. TPCK-Trypsin was added for a final concentration of 1ug/mL, and incubated for 4 hours, 10^5^ MDCK cells were then added to the well, and supernatant taken every 24 hours and assayed by TCID_50_ to determine viral load. Virus was plaque purified and was confirmed to grow in the presence of 23mM oseltamivir carboxylate (SOURCE), and not to grow at 37 degrees. Stocks for further assay were grown in MDCK cells infected at an MOI∼0.01 in Minimal Essential Media (Gibco) supplemented with: BSA fraction V (Gibco), GlutaMax (Gibco), 1ug/mL TPCK-Trypsin (ThermoScientific 20233), 23mM oseltamivir carboxylate at 33 degrees in a 5% CO_2_ atmosphere. After 96 hours of infection and complete destruction of the monolayer, supernatant was harvested, centrifuged at 5000 xg to pellet debris and was aliquoted and stored at −70C. Titer was determined by 50% tissue culture infectious dose (TCID_50_).

### Determination of TCID_50_

Viral titer was determined by 50 percent tissue culture infectious dose assay. MDCK cells were seeded at 3×10^4^ cells per well in 96 well plates (Costar). Serial 10 fold dilutions of viral samples were made in Infection media. Cells were infected in triplicate with each dilution. 72 hours post infection, supernatant from infected cells was mixed 1:1 with 0.5% turkey red blood cells in PBS (Lampire) in V bottom 96 well plates for hemagglutination assay. Plates were incubated at room temperature for 30 minutes. Hemagglutination was read for each well and 50% tissue culture infectious dose was calculated using the method of Reed and Munch(52).

### Mouse Infection

All experiments involving animals were performed with prior approval of and in accordance with guidelines of the St. Jude Institutional Animal Care and Use Committee. The St. Jude laboratory animal facilities are fully accredited by the American Association for Accreditation of Laboratory Animal Care. Laboratory animals are maintained in accordance with the applicable portions of the Animal Welfare Act and the guidelines prescribed in the DHHS publication *Guide for the Care and Use of Laboratory Animals.* 6-8 week old female C57/Bl6 mice were purchased from Jackson labs and acclimated in the facility at least one week prior to infection. Mice were treated with 25mg/kg oseltamivir phosphate (Sigma Aldrich) by oral gavage two hours prior to viral infection and every twelve hours post viral challenge. Mice were infected under 3% isoflorane sedation by the intranasal route with 3 log TCID_50_ wild type PR8, 3 log TCID_50_ temperature sensitive/oseltamivir resistant PR8, a mix of the two viruses, or a mix of the two viruses pre-mixed with either live or ethanol fixed *S. pneumoniae* for a final concentration of 3 log TCID_50_ of virus and 10^6^ CFU of bacteria in a volume of 50uL. Mice were weighed daily for four days to determine signs of severe viral illness. At 4 days post infection, lungs were harvested and homogenized in 0.5mL phosphate buffered saline and both total viral load and oseltamivir resistant temperature resistant viral load was determined by TCID_50_ in the presence and absence of 23uM oseltamivir carboxylate (Sigma Aldrich).

### Reassortment in A549 and Detroit 562 cells

A549 and Detroit 562 cells were seeded into 24 well tissue culture plates (Costar) at a density of 5×10^4^ cells/well in a volume of 0.5mL per well 24 hours prior to viral infection. Bacteria were grown to mid-logarithmic growth phase and were normalized to 10^8^ CFU/mL in viral infection media without trypsin. Inocula were generated with 4 log TCID_50_/mL wild type and/or temperature sensitive PR8 and 10^5-7^ CFU/mL live or ethanol killed bacteria in viral infection media without trypsin. Inocula were mixed by rocking at 37 degrees. Cells were washed twice with phosphate buffered saline and infected with 0.5mL viral inoculum for one hour at 33 degrees C in a 5% CO_2_ atmosphere. After 1 hour, inoculum was removed and replaced with viral infection media without trypsin and supplemented with PenStrep (Gibco) (final concentration 100 U/mL Penicillin, 100 U/mL Streptomycin) and cells incubated at permissive temperature, 33 degrees in a humidified 5% CO_2_ atmosphere for 20 hours to allow one round of viral replication. Supernatants were collected and stored at −70C until TCID_50_ was determined at non-permissive temperature (37 C) and in the presence of 23uM oseltamivir carboxylate to determine the output of reassortant virus. 3-9 wells were infected total per group over 3 independent experiments.

#### Catalase treatment

To determine the impact of bacterial and spontaneous hydrogen peroxide production, 0.1% catalase (MP Biomedicals) or equal volume of water for control experiments was added to 4 log TCID_50_/mL wild type and/or temperature sensitive PR8 and 10^6^ CFU/mL live bacteria in viral infection media without trypsin and incubated with rocking at 37 degrees. Cells were washed twice with phosphate buffered saline and infected with 0.5mL viral inoculum. After 1 hour, inoculum was removed, and replaced with viral infection media without trypsin and supplemented with PenStrep and 0.1% catalase or equal volume of water as control, and infection proceeded as above.

#### TLR4 inhibitor treatment

TAK-242 (Sigma Aldrich) was resuspended in dimethyl sulfoxide (DMSO) (Sigma Aldrich). A549 and Detroit 562 cells were seeded into 24 well tissue culture plates (Costar) at a density of 5×10^4^ cells/well in a volume of 0.5mL per well in media supplemented with 1uM TAK-242, or 0.004% DMSO as vehicle control 24 hours prior to viral infection. TAK242 or DMSO was maintained through inoculum generation and viral infection.

### Statistical Analyses

Statistical analyses were performed using GraphPad Prism Version 10.6.1. Data was tested for normality with the Kolmogorov-Smirnov test. If data were normal, unpaired t-tests were performed, if data were non-normal, the Mann-Whitney test was performed.

## Acknowledgements

HMR received funding from Medical Research Foundation of Oregon New Investigator Award 2022 and startup funding from Oregon State University. Stacey Schultz-Cherry and Jason Rosch at St Jude Children’s Research Hospital provided mentorship during the development of this project and ALSAC provided funds.

---

**Supplemental Figure 1:**
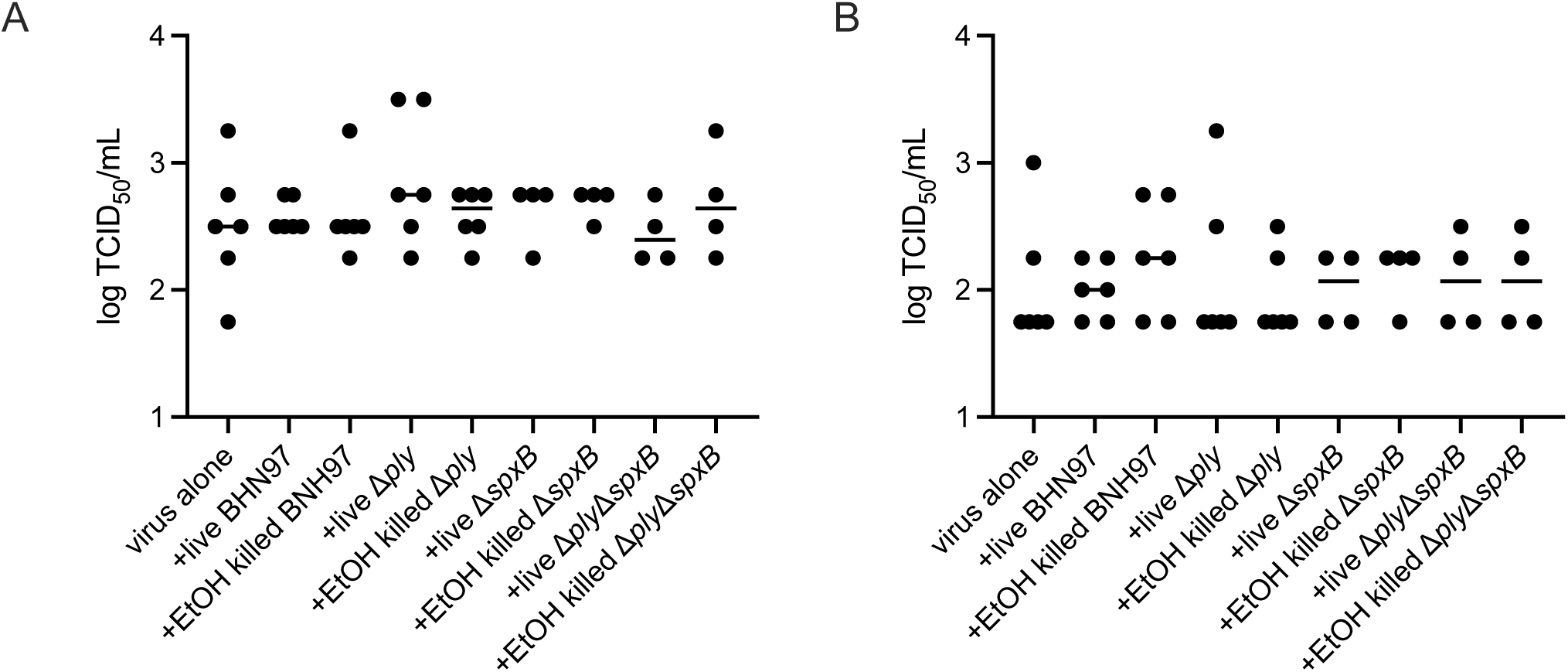
Viability and toxin production by *S. pneumoniae* does not impact viral replication. **A)** A549 or **B)** Detroit-562 cells were infected with IAV strain PR8 pre-incubated with the indicated bacterial strain. Following one hour of infection, inocula were removed and replaced with infection media containing antibiotics. 24 hours post infection, supernatants were collected and viral replication was determined by TCID_50_ on MDCK cells. Data are presented as titer from each cell culture well as a single data point with the bar representing the median.

**Supplemental Figure 2:**
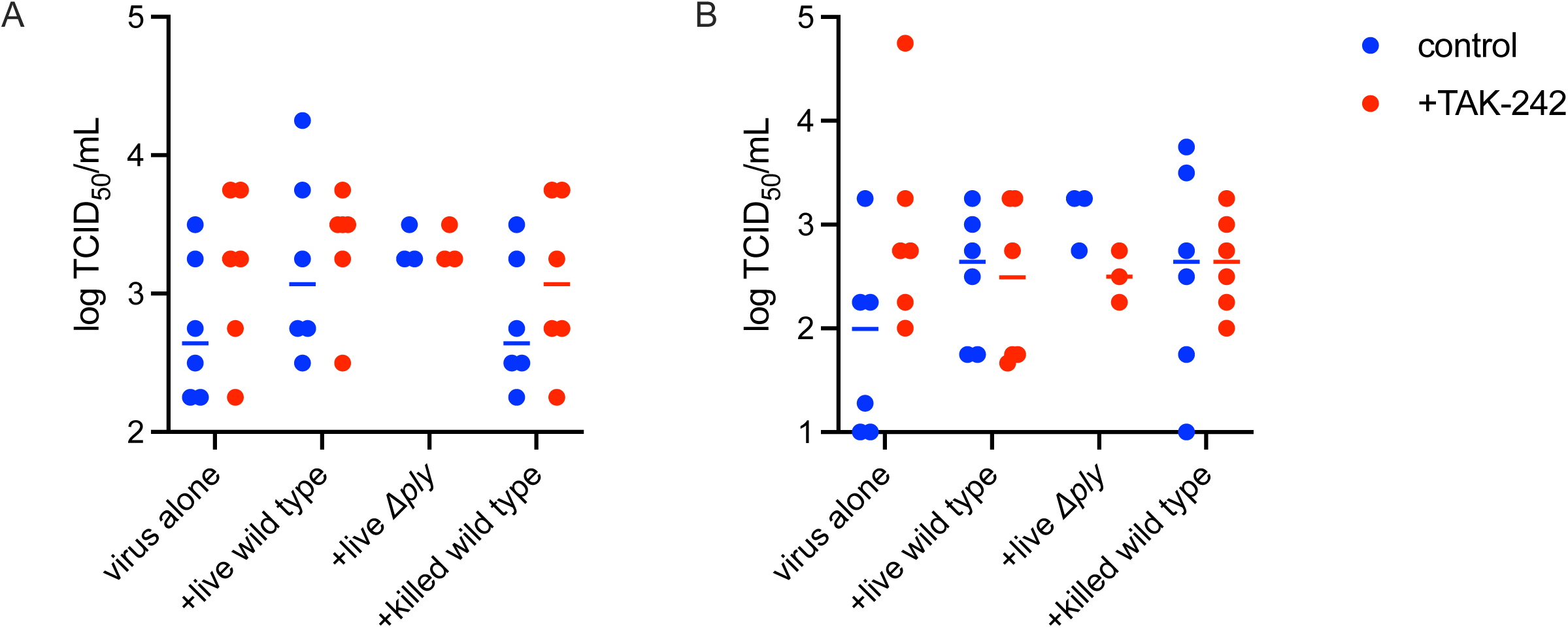
TLR4 inhibition with or without co-infection with *S. pneumoniae* does not impact viral replication. **A)** A549 or **B)** Detroit-562 cells were pre-treated with, inocula were prepared in, and cells were infected in the presence of 1μM TAK-242, or 0.004% DMSO vehicle control. Cells were infected with IAV strain PR8 pre-incubated with the indicated bacterial strain. Following one hour of infection, inocula were removed and replaced with infection media containing antibiotics. 24 hours post infection, supernatants were collected and viral replication was determined by TCID_50_ on MDCK cells. Data are presented as titer from each cell culture well as a single data point with the bar representing the median.

